# Ancient genomes from North Africa evidence prehistoric migrations to the Maghreb from both the Levant and Europe

**DOI:** 10.1101/191569

**Authors:** Rosa Fregel, Fernando L. Méndez, Youssef Bokbot, Dimas Martín-Socas, María D. Camalich-Massieu, Jonathan Santana, Jacob Morales, María C. Ávila-Arcos, Peter A. Underhill, Beth Shapiro, Genevieve Wojcik, Morten Rasmussen, Andre E. R. Soares, Joshua Kapp, Alexandra Sockell, Francisco J. Rodríguez-Santos, Abdeslam Mikdad, Aioze Trujillo-Mederos, Carlos D. Bustamante

**Affiliations:** Department of Genetics, School of Medicine, Stanford University, 365 Lasuen Street, CA 94305, Stanford, US.; Institut National des Sciences de l’Archéologie et du Patrimoine, Avenue Allal el Fassi, 6828, Rabat, Morocco.; Departmento de Prehistoria, Universidad de La Laguna, Calle Prof. Jose Luis Moreno Becerra, 38320, San Cristóbal de La Laguna, Spain.; Department of Archaeology, Durham University, South Rd, Durham DH1 3LE, United Kingdom.; Departamento de Ciencias Históricas, Universidad de Las Palmas de Gran Canaria, Pérez del Toro 1, 35003, Las Palmas de Gran Canaria, Spain.; International Laboratory for Human Genome Research, National Autonomous University of Mexico, Blvd. Juriquilla 3001, 76230, Querétaro, Mexico.; Department of Ecology and Evolutionary Biology, University of California, 1156 High Street, CA 95064, Santa Cruz, US.; Instituto Internacional de Investigaciones Prehistóricas de Cantabria, Avda. de los Castros, 39005, Cantabria, Spain.

**Keywords:** ancient DNA, paleogenomics, Neolithic, North Africa

## Abstract

The extent to which prehistoric migrations of farmers influenced the genetic pool of western North Africans remains unclear. Archaeological evidence suggests the Neolithization process may have happened through the adoption of innovations by local Epipaleolithic communities, or by demic diffusion from the Eastern Mediterranean shores or Iberia. Here, we present the first analysis of individuals’ genome sequences from early and late Neolithic sites in Morocco, as well as Early Neolithic individuals from southern Iberia. We show that Early Neolithic Moroccans are distinct from any other reported ancient individuals and possess an endemic element retained in present-day Maghrebi populations, confirming a long-term genetic continuity in the region. Among ancient populations, Early Neolithic Moroccans are distantly related to Levantine Natufian hunter-gatherers (∼9,000 BCE) and Pre-Pottery Neolithic farmers (∼6,500 BCE). Although an expansion in Early Neolithic times is also plausible, the high divergence observed in Early Neolithic Moroccans suggests a long-term isolation and an early arrival in North Africa for this population. This scenario is consistent with early Neolithic traditions in North Africa deriving from Epipaleolithic communities who adopted certain innovations from neighbouring populations. Late Neolithic (∼3,000 BCE) Moroccans, in contrast, share an Iberian component, supporting theories of trans-Gibraltar gene flow. Finally, the southern Iberian Early Neolithic samples share the same genetic composition as the Cardial Mediterranean Neolithic culture that reached Iberia ∼5,500 BCE. The cultural and genetic similarities of the Iberian Neolithic cultures with that of North African Neolithic sites further reinforce the model of an Iberian migration into the Maghreb.

**SIGNIFICANCE STATEMENT:** The acquisition of agricultural techniques during the so-called Neolithic revolution has been one of the major steps forward in human history. Using next-generation sequencing and ancient DNA techniques, we directly test if Neolithization in North Africa occurred through the transmission of ideas or by demic diffusion. We show that Early Neolithic Moroccans are composed of an endemic Maghrebi element still retained in present-day North African populations and distantly related to Epipaleolithic communities from the Levant. However, late Neolithic individuals from North Africa are admixed, with a North African and a European component. Our results support the idea that the Neolithization of North Africa might have involved both the development of Epipaleolithic communities and the migration of people from Europe.

## INTRODUCTION

One of the greatest transitions in human history was the transition from hunter-gatherer to farming lifestyle. How farming traditions expanded from their birthplace in the Fertile Crescent has been a matter of contention. Two models were proposed: one involving the movement of people and the other based on the transmission of ideas. Over the last decade, paleogenomics has been instrumental in settling long-disputed archaeological questions^1^, including those surrounding the Neolithic revolution^2^. Compared to the extensive genetic work done on Europe and the Near East, the Neolithic transition in North Africa, including the Maghreb, remains largely uncharacterized. Archaeological evidence suggests that some of the major innovations associated with the Neolithic, such as farming and pottery production, could have been introduced into northern Morocco through sea voyaging by people from Iberia or the central Mediterranean as early as *ca*. 5400 BCE^3,4^. In fact, some of the Neolithic pottery recorded in North Africa strongly resembles that of European cultures like the Cardial Early Neolithic, the Mediterranean early farmer culture located in Iberia^5^. However, other innovations such as some pottery traditions and bone and lithic technical customs could be the result of *in situ* development from Epipaleolithic communities, indicating a strong continuity in the local population since the Late Pleistocene^6-10^.

Genetic data from present-day populations^11-13^ suggests that North African ancestry has contributions from four main sources: 1) an autochthonous Maghrebi component related to a back migration to Africa ∼12,000 years ago from Eurasia; 2) a Middle Eastern component probably associated with the Arab conquest; 3) a sub-Saharan component derived from trans-Saharan migrations; and 4) a European component that has been linked to recent historic movements. Paleogenomic studies have begun to provide insights into North African Prehistory^14-16^; however, no research to date has tested whether the Neolithic transition in the Maghreb was driven by local populations who adopted cultural and technological innovations or the migration of people. Here, we perform genome-wide analysis of remains from the Early Neolithic site of Ifri n’Amr or Moussa (IAM; ∼5,200 BCE, n=7) and the Late Neolithic site of Kelif el Boroud (KEB; ∼3,000 BCE; n=8) (Supplementary Note 1). To test possible migrations through the Strait of Gibraltar, we also analyse human remains from the southern Iberian Early Neolithic site of El Toro (TOR; ∼5,000 BCE; n=12) (Figure 1). This Iberian Early Neolithic culture bears similarities with early Maghrebi pottery decoration, as well as bone and lithic tool production traditions which suggest an African influence^17^ (Supplementary Note 1). Including these southern Iberian samples in our analysis enables a direct test of this hypothesis.

**Figure 1.**
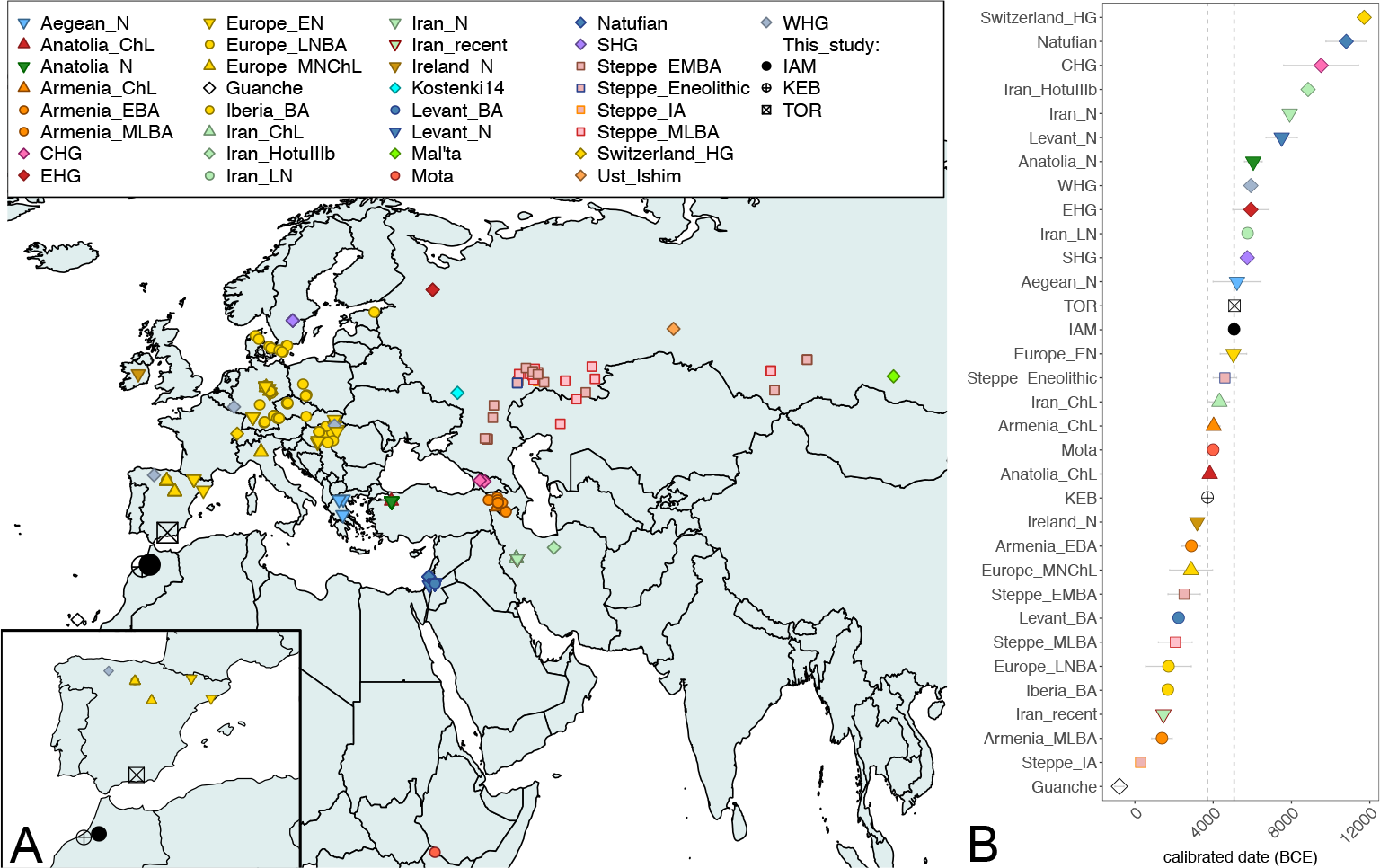
Geographical location (A) and calibrated radiocarbon date (B) of the samples included in this study, as well as other ancient DNA samples from the literature.

## RESULTS AND DISCUSSION

We sequenced 38 Illumina pair-end libraries from 27 individuals, and selected the best-conserved libraries for subsequent analyses. Endogenous DNA content was generally low (2.88% on average) (Supplementary Note 2). Depth of coverage was consistently improved when enriching using baits targeting specific sites for the Multiethnic Genotyping Array (MEGA) (∼100X), compared to whole-genome capture (∼15X) (Supplementary Note 2). Following enrichment, we generated thirteen low-coverage genomes (five from IAM, four from KEB and four from TOR), with MEGA coverage ranging from 0.04X to 1.72X depth, and genome-wide coverage ranging from 0.01X to 0.74X depth (Table 1). All samples considered in this study met the standard aDNA authentication criteria, including observation of DNA fragmentation (∼46 bp average read length) and damage patterns due to cytosine deamination toward the 5′ ends of molecules (Supplementary Note 3).

**Table 1.**
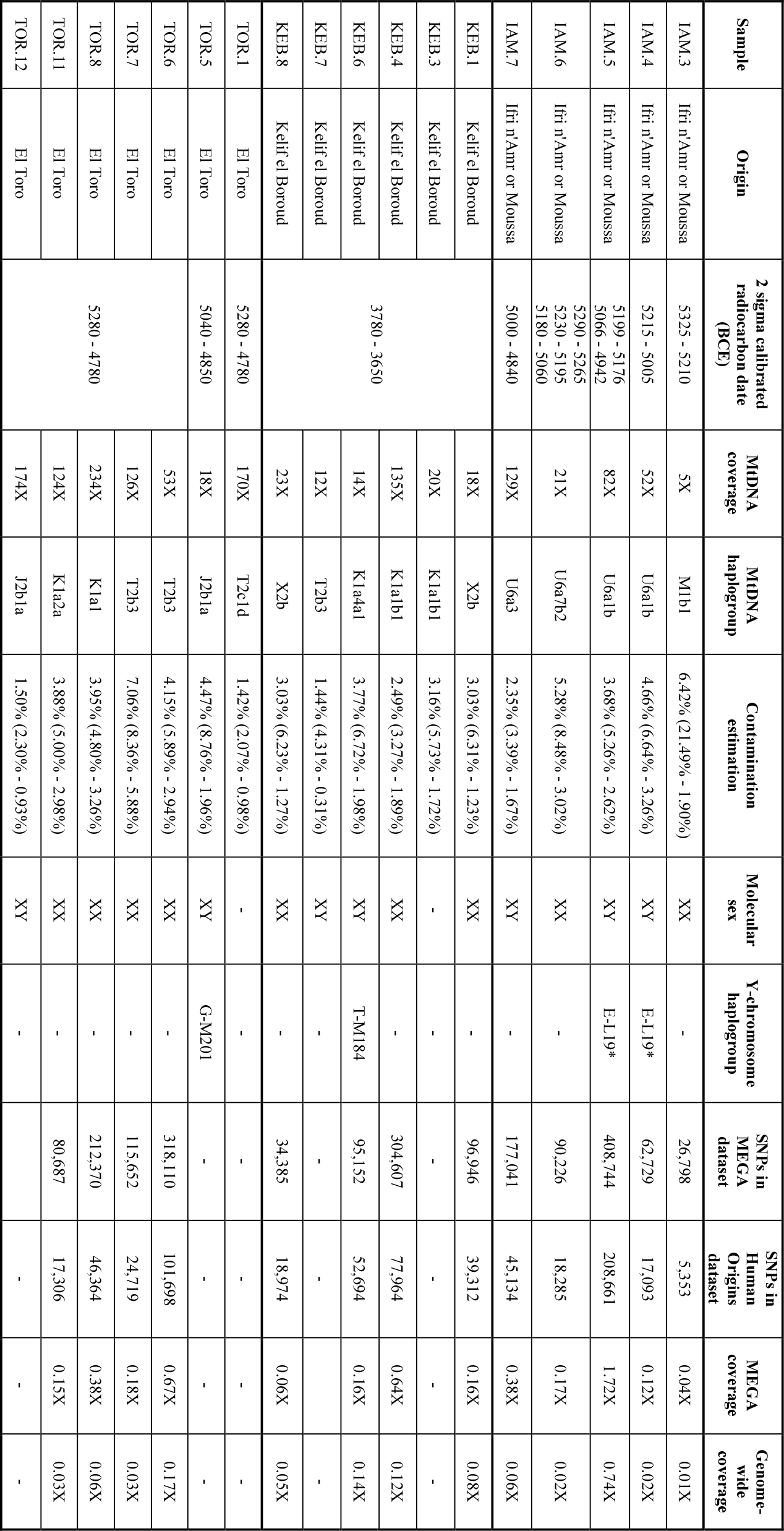
Summary statistics for North African and Iberian samples.

Mitochondrial DNA and Y-chromosome haplogroups obtained for IAM (Moroccan Early Neolithic) and KEB (Moroccan Late Neolithic) suggest either a population replacement or an important genetic influx into Morocco between 5,000–3,000 BCE. IAM samples belong to the mtDNA haplogroups U6a and M1—both of which are associated with the back migration to Africa from Eurasia in Upper Palaeolithic times^18,19^—while KEB samples belong to haplogroups K1, T2 and X2, prominently found in Anatolian and European Neolithic samples^2,20^ (Supplementary Note 4). Regarding the paternal lineages, IAM individuals carry Y chromosomes distantly related to the typically North African E-M81 haplogroup, while the Y chromosome from KEB belongs to the T-M184 haplogroup; though scarce and broadly distributed today, this haplogroup has also been observed in European Neolithic individuals^16^ (Supplementary Note 5). Both mtDNA and Y-chromosome lineages (K1, J2 and T2 haplogroups, and G-M201 haplogroup, respectively) for samples from TOR (Iberian Early Neolithic) are similar to those observed in Europe during Neolithic times^20^.

When projected on a Principal Components Analysis (PCA) space built using sub-Saharan African, North African, European and Middle Eastern population of the Human Genome Diversity Project (HGDP) dataset genotyped with MEGA, IAM samples are placed close to Mozabites, while Iberian Neolithic samples fall close to southern European populations (Supplementary Note 6). As suspected from the mtDNA and Y-chromosome data, KEB samples do not cluster with IAM and are placed in an intermediate position between IAM and TOR. We further explored the genetic structure of these samples using the program ADMIXTURE^21^ (Figure 2). At K=5, we observe sub-Saharan African (red), early European Neolithic (green), North African (yellow), Middle Eastern (violet) and eastern European components (orange). Congruently with PCA results, TOR is composed of the early European component, clustering with Sardinian samples, and IAM is composed of the North African component, clustering with Mozabites. Finally, KEB is placed in an intermediate position, with ∼50% of both early European Neolithic and North African ancestries. It is worth mentioning that, compared to current North African samples, IAM and KEB do not show any sub-Saharan African ancestry, suggesting that trans-Saharan migrations occurred after Neolithic times. This is in agreement with the analysis of present-day genome-wide data from Morocco, which estimated a migration of western African origin into Morocco only ∼1,200 years ago^11^.

**Figure 2.**
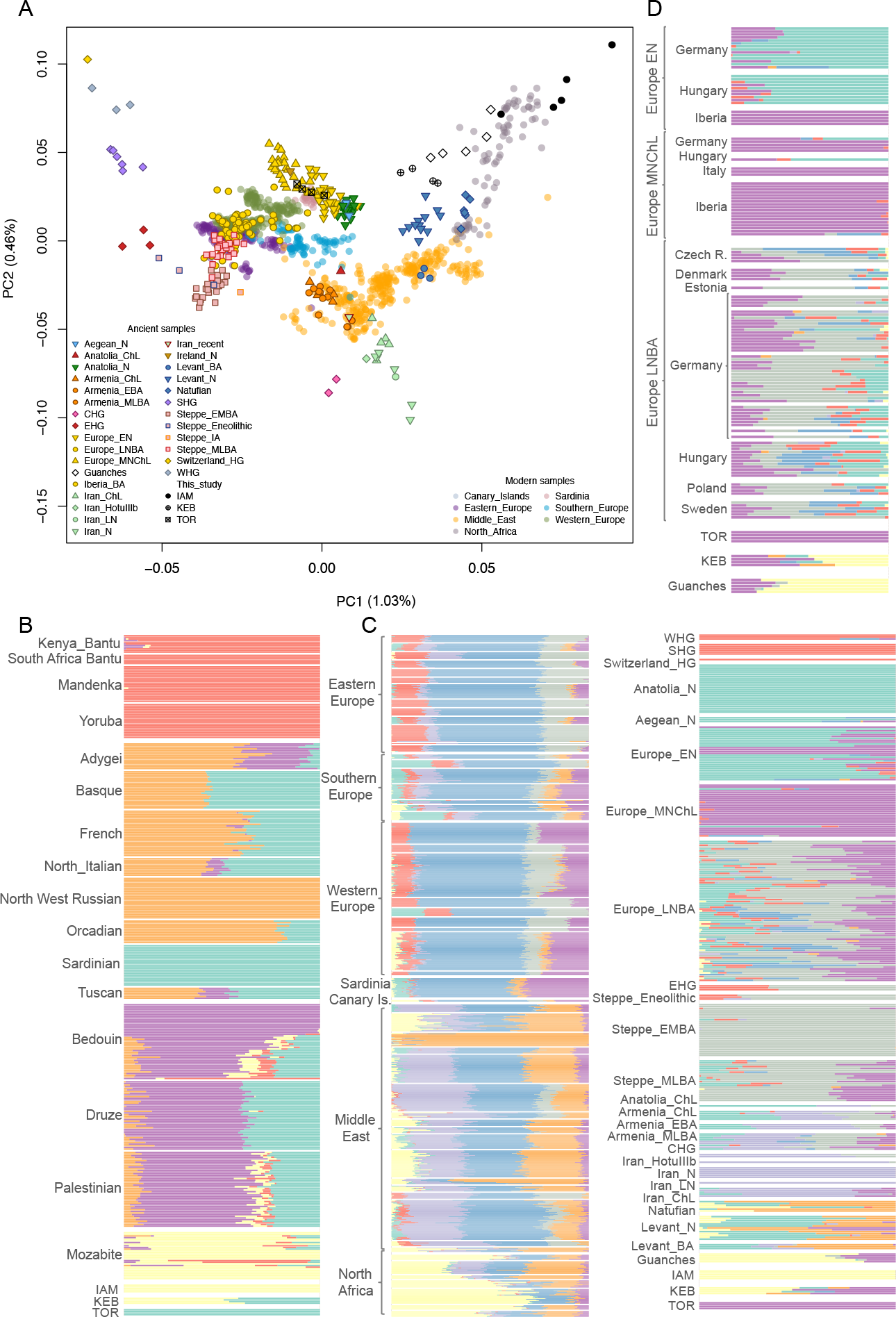
Ancestry inference in ancient samples from North Africa and the Iberian Peninsula. (A) PCA analysis using the Human Origins panel, (B) ADMIXTURE analysis using the HGDP-MEGA dataset (K=5), (C) ADMIXTURE analysis using the Human Origins dataset for modem and ancient populations (K=8), and (D) detailed ADMIXTURE analysis for European Neolithic samples (K=8).

West Eurasian populations can be modelled as admixture of four different ancestral components^2^: Eastern and Western European hunter-gatherers, and Iranian and Levantine Neolithic. We explored the placement of Moroccan and Southern Iberian Neolithic samples in this context, and compared their genetic affinities to ancient and present-day West Eurasian and Levant populations in the Human Origins panel. Interestingly, PCA reveals that IAM individuals are different from any aDNA sample studied to date (Figure 2; Supplementary Note 6). When projected, IAM samples are close to modern North Africans, in the Levantine corner of the PCA space (Figure 2). Southern Iberian Neolithic individuals from TOR cluster with Sardinians and with other Anatolian and European Neolithic samples. Moreover, KEB samples are placed halfway between the IAM and Anatolian/European farmer clusters, in close proximity to Levant aDNA samples and also to Guanche samples^16^, the indigenous population of the Canary Islands known to have a Berber origin^22^. When compared using ADMIXTURE (See Supplementary Note 7 for details), IAM samples possess ∼100% of a component partially shared by aDNA samples from the Middle East and Levant at low K values. At K=6, this IAM-like component is observed mainly in modern North Africa, following a west-to-east cline, and in the Guanches. TOR and other Early Neolithic samples from Iberia cluster together with farmers from Anatolia, the Aegean area and Europe. At K=8, the Early Neolithic individuals from Iberia differentiate from the Anatolian, Aegean and European Early Neolithic samples, and share their main component (purple) with Middle Neolithic/Chalcolitic samples (Figure 2). Finally, at low K values, KEB can be explained as having both IAM-like and European Neolithic components, suggesting an admixture process between IAM-like people and early farmers. Nevertheless, at K=8, the European component in KEB is predominantly “purple,” with some “green” component. This “green” component is also present, at a low frequency, in Natufians and other ancient Levantine populations. The substantially larger contribution of the “purple” component, when compared with the “green”, suggests a significant genetic contribution of ancient Iberians in Morocco (Figure 2). The same admixture profile is observed in Guanches, but the amount of IAM ancestry is consistently higher in all the samples. Given that the Guanches could have had originated in a different area of the Maghreb, this result might suggest that the European Neolithic impact in North Africa was heterogeneous.

To compare our samples directly to the genomes of ancient and modern populations, we calculated pair-wise FST distances, which, unlike PCA and global ancestry analyses, are insensitive to the inclusion of large numbers of individuals from modern populations. FST values indicate that the IAM samples are as differentiated from all other populations as Yoruba are from non-Africans (Supplementary Note 9), with the sole exception of KEB and, to a lesser extent, the Guanches and modern North African populations. Given the relatively low heterozygosity and high identity-by-descent proportions observed in IAM (Supplementary Note 8), this differentiation could be driven by isolation and genetic drift. IAM is divergent from the other populations, with the exception of populations that likely received genetic influx from them. This raises the possibility that IAM was isolated in North Africa since Palaeolithic times, when a back migration from Eurasia brought mtDNA haplogroups M1 and U6 to the Maghreb^18^. Although IAM is clearly more similar to KEB than to any other population, the converse is not true. KEB has lower F_ST_ distances with any Anatolian, European (excluding European hunter gatherers), Levantine and Iranian population, rather than with IAM. In the modern DNA reference panel, KEB is similar to North African, European and Middle Eastern populations. Among the ancient populations, TOR is more similar to Middle Neolithic/Chalcolithic Europeans, and, among modern populations, to populations from Spain, North Italy and Sardinia.

To further investigate the genetic affinities of IAM, KEB and TOR samples, we conducted outgroup f3-statistic analysis^23^. Results indicate that, when KEB and Guanches are excluded, IAM shares more drift with ancient Levantine populations, such as Natufians (Epipaleolithic) and Pre-Pottery Neolithic individuals (Figure 3; Supplementary Note 10), than with any other ancient population. To explore further the connection between IAM and Levantine populations, we performed an f4-statistic analysis to test whether IAM shares more alleles with any other population in the Human Origins panel^2,24^ than with ancient populations from the Levant (Supplementary Note 10). Consistently, and also with the exception of KEB and Guanches, all comparisons indicated higher similarity with Natufians and Levantine farmers. This suggests that most of IAM ancestry originates from an out-of-Africa source, as IAM shares more alleles with Levantines than with any sub-Saharan Africans, including the 4,500-year-old genome from Ethiopia^14^. To further test the hypothesis that IAM is more closely related to out-of-Africa populations, we determined if we could detect Neanderthal ancestry in IAM, which is typical of non-African populations. A signal of Neanderthal ancestry has been detected in modern North African populations^25^. A lack of Neanderthal ancestry in IAM would imply that the signal observed today is a product of more recent migration into North Africa from the Middle East and Europe in historical times. When compared to the Neanderthal high coverage genome sequence from Altai^26^ and the low-coverage sequence from Vindija Cave^27^, and using the S-statistic^23^, we detected a Neanderthal introgression signal into IAM, suggesting derivation from the same event shared by non-African populations. All these results together indicate that the origin of IAM was outside Africa, most probably from the Levant. However, it is important to take into account that the number of ancient genomes for comparison is low and future sampling can provide further refinement in the origin of IAM.

**Figure 3.**
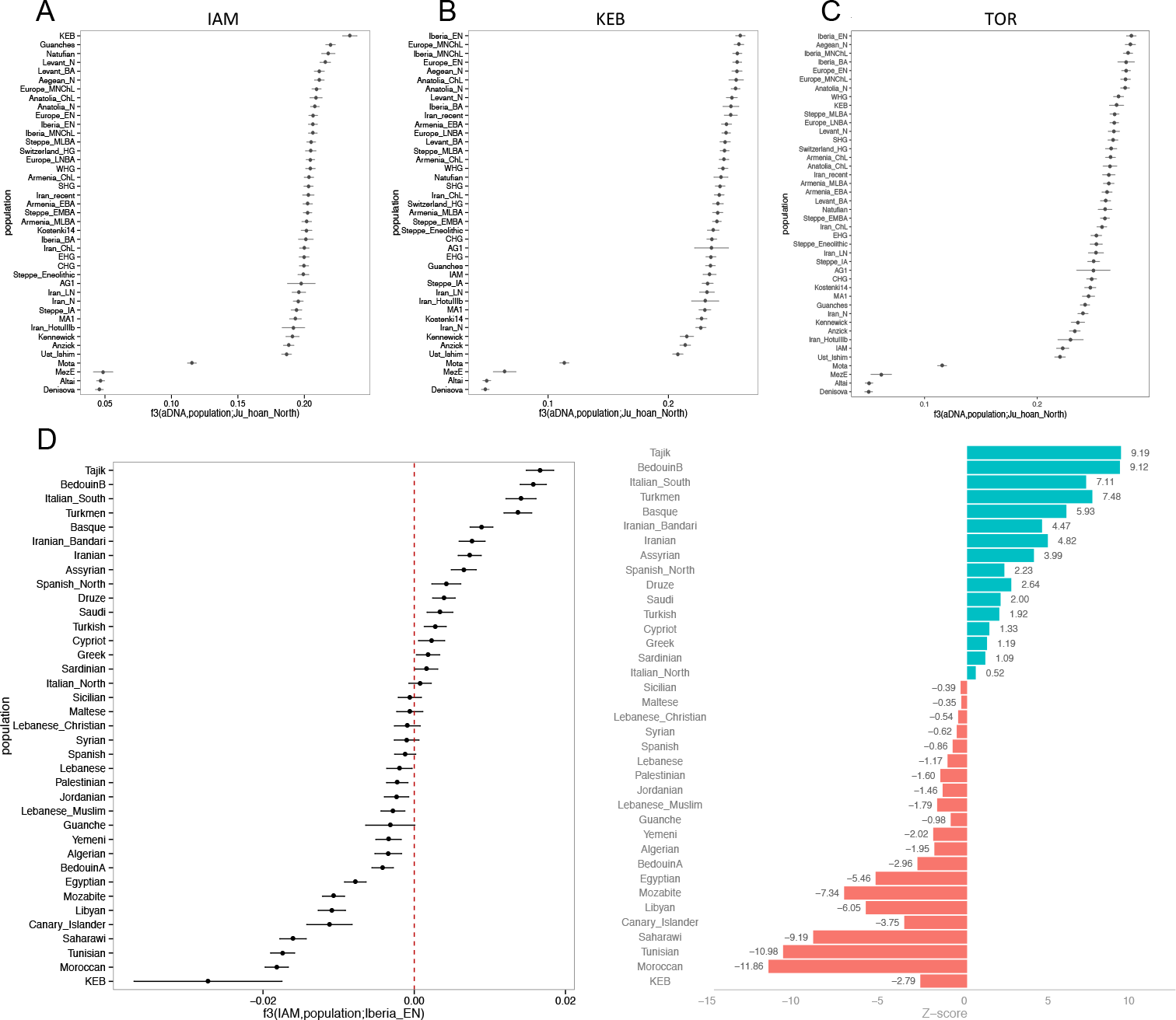
Outgroup f3-statistic for IAM (A), KEB (B) and TOR (C), and admixture f3-statistic for KEB (D).

Both F_ST_ and outgroup-f3 statistic analyses indicate that KEB shares ancestry with IAM, but also more genetic drift with Neolithic and Chalcolithic populations from Anatolia and Europe, with the highest shared genetic drift appearing in Iberian Early Neolithic samples (Figure 3; Supplementary Note 10). This pattern and the result from ADMIXTURE could be explained if the KEB population was a mixture between IAM-related and European Neolithic groups. To formally test this hypothesis, we used an admixture-f3 test^23^, using KEB as the test population, IAM as a reference population and one of the Anatolian and European Neolithic and Chalcolithic populations as the second reference population. All comparisons produced negative values of the f3-statistic, which suggests the KEB population can be modelled as a mixture of IAM and Anatolian/European Neolithic.

TOR has more shared ancestry with Iberian Early Neolithic samples and other Neolithic and Chalcolithic populations from Europe. Archaeological studies have suggested that there was an Andalusian Early Neolithic culture with North African influences before the Cardial expansion into the Western Mediterranean basin^28^. However, we observe that TOR samples have a similar genetic composition to that of Cardial individuals from Iberia, evidencing a common origin, and ruling out an Andalusian Early Neolithic distinct from Cardial Culture.

Finally, although limited by low coverage, phenotypic predictions based on genetic variants of known effects agree with our estimates of global ancestry. IAM people do not possess any of the European SNPs associated with light pigmentation, and most likely had dark skin and eyes. IAM samples present ancestral alleles for pigmentation-associated variants present in SLC24A5 (rs1426654), SLC45A2 (rs16891982) and OCA2 (rs1800401 and 12913832) genes. On the other hand, KEB individuals exhibit some European-derived alleles that predispose individuals to lighter skin and eye colour, including those on genes SLC24A5 (rs1426654) and OCA2 (rs1800401) (Supplementary Note 11).

## CONCLUSION

Genetic analyses have revealed that the population history of modern North Africans is quite complex^11^. Based on our aDNA analysis, we identify an Early Neolithic Moroccan component that is restricted to North Africa in present-day populations^11^, and that is the sole ancestry in IAM samples. We hypothesize that this component represents the autochthonous Maghrebi ancestry associated with Berber populations. This Maghrebi component is different from those of any ancient samples studied so far and is distantly related to that of Epipaleolithic people from the Levant. Our data suggests that the IAM population was isolated in the Maghreb since the Upper Palaeolithic back migration, although it is impossible to be certain without paleogenomic data from North African Palaeolithic samples. An expansion in Early Neolithic times followed by strong genetic drift might also be plausible.

Our hypothesis is in agreement with archaeological research pointing to the first stage of the Neolithic expansion in Morocco as the result of a local population who adopted some technological innovations, such as pottery production or farming, from neighbouring areas. By 3,000 BCE, a continuity in the Neolithic spread brought Mediterranean-like ancestry to the Maghreb, most likely from Iberia. Other archaeological remains, such as African elephant ivory and ostrich eggs found in Iberian sites, confirm the existence of contacts and exchange networks through both sides of the Gibraltar strait at this time. Our analyses strongly support that at least some of the European ancestry observed today in North Africa is related to prehistoric migrations, and local Berber populations were already admixed with Europeans before the Roman conquest. Furthermore, additional European/Iberian ancestry could have reached the Maghreb after KEB people; this scenario is supported by the presence of Iberian-like Bell-Beaker pottery in more recent stratigraphic layers of IAM and KEB caves. Future palaeogenomic efforts in North Africa will further disentangle the complex history of migrations that forged the ancestry of the admixed populations we observe today.

## MATERIAL AND METHODS

Measures to avoid and monitor contamination from modern DNA were applied, at all times, during sample manipulation. Ancient DNA was extracted from teeth or bone, built into double-stranded indexed libraries and sequenced on an Illumina NextSeq 500 (Supplementary Note 2). Due to the environmental conditions of the burial sites, we expected to recover low proportions of endogenous DNA from these ancient remains. To overcome limitations due to DNA degradation, we applied two different capture methods to enrich for human reads (Supplementary Note 2): one targeting the whole genome^29^ and one targeting the variants of the MEGA array (Illumina Inc.).

Reads were trimmed and adapters removed using AdapterRemoval^30^, and then mapped to the human reference genome (hg19) using BWA^31^. Low quality (MAPQ<30) and duplicate reads were removed using SAMtools^32^ MapDamage^33^ was used to visualize misincorporation and fragmentation patterns, and to rescale the quality of bases likely affected by post-mortem damage. Confidence intervals of sex determination were calculated following Skoglund et al.^34^. MtDNA haplogroups were determined using HaploGrep^35^. Y-chromosome haplogroup inference was carried out as in Schroeder et al.^36^. As the reference panel, we used both the Human Origins panel^2^ and the HGDP dataset genotyped with MEGA-ex (Illumina Inc.). For principal component analysis, we projected the aDNA samples on the PCA space built with the modern dataset, using smartpca^37^ and LASER^38^. Admixture estimations were done using ADMIXTURE software^21^. F_ST_ distances were calculated using smartpca^37^. Identity-by-descent proportions were estimated using PLINK^39^, and heterozygosity estimations using a newly developed method for low-coverage genomes (Supplementary Note 8). f-statistics estimates were calculated using admixtools software^23^. All plots were prepared using R software^40^. Detailed information about methods is included in the Supplementary Notes.

## Acknowledgements

C.D.B and R.F. were funded by a grant from the National Science Foundation (1201234). R.F. was funded by Fundación Canaria Dr. Manuel Morales fellowship. A.E.R.S. was funded by Ciencia sem Fronteiras fellowship - CAPES, Brazil. B.S. and J.K. were funded by a grant from the Gordon and Betty Moore Foundation (GBMF-3804).

## Author Contributions

R.F. and C.D.B conceived the idea for the study, D.M.S, M.D.C.M, J.S., J.M, Y.B., F.J.R.S, A.M. and A.T.M assembled skeletal material and provided archaeological background for the samples, R.F., A.E.R.S., J.K. and A.S. performed work in the wet laboratory, F.M. developed methods for data analysis, M.R. developed methods for DNA capture, R.F., F.M., M.A.A, P.A.U, G.W., analysed data, C.D.B and B.S. supervised the study, R.F., F.M. and C.D.B. wrote the manuscript and supplements with input from all co-authors.

## Author Information

Sequence data are available through the European Nucleotide Archive (PRJEB22699). Consensus mtDNA sequences are available at the National Center of Biotechnology Information (Accession Numbers MF991431-MF991448). The authors declare no competing financial interests. Correspondence and requests for materials should be addressed to R.F.

## REFERENCES

1 Nielsen, R. et al. Tracing the peopling of the world through genomics. Nature 541, 302-310, doi: 10.1038/nature21347 (2017).

2 Lazaridis, I. et al. Genomic insights into the origin of farming in the ancient Near East. Nature 536, 419-424, doi:10.1038/nature19310 (2016).

3 Linstädter, J., Medved, I., Solich, M. & Weniger, G. C. Neolithisation process within the Alboran territory: Models and possible African impact. Quaternary International 274, 219 - 232, doi:10.1016/j.quaint.2012.01.013 (2012).

4 Zilhão, J. Early prehistoric navigation in the Western Mediterranean: Implications for the Neolithic transition in Iberia and the Maghreb. Eurasian Prehistory, 185-200(2014).

5 Martínez-Sánchez, R. M., Vera-Rodríguez, J. C., Pérez-Jordá,G., Peña-Chocarro, L. & Bokbot, Y. The beginning of the Neolithic in northwestern Morocco. Quaternary International In press (2017).

6 Mulazzani, S. et al. The emergence of the Neolithic in North Africa: A new model for the Eastern Maghreb. Quaternary International 410, 123 - 143, doi:10.1016/j.quaint.2015.11.089 (2016).

7 Barton, N. et al. Human Burial Evidence from Hattab II Cave and the Question of Continuity in Late Pleistocene–Holocene Mortuary Practices in Northwest Africa. Cambridge Archaeological Journal 18, 195–214, doi:10.1017/S0959774308000255 (2008).

8 Lubell, D., Sheppard, P. & Jackes, M. in Advances in World Archaeology (eds F. Wendorf & A. Close) 146-191 (Academic Press, 1984).

9 Linstädter, J. Epipalaeolithic-Neolithic-Transition in the Mediterranean region of Northwest-Africa. Quartär 55, 41-62 (2008).

10 de Groote, I. & Humphrey, L. T. Characterizing evulsion in the Later Stone Age Maghreb: Age, sex and effects on mastication. Quaternary International 413, 50 - 61, doi:https://doi.org/10.1016/j.quaint.2015.08.082 (2016).

11 Henn, B. M. et al. Genomic Ancestry of North Africans Supports Back-to-Africa Migrations. Plos Genetics 8, e1002397-e1002397, doi:10.1371/journal.pgen.1002397 (2012).

12 Arauna, L. R. et al. Recent Historical Migrations Have Shaped the Gene Pool of Arabs and Berbers in North Africa. Mol Biol Evol 34, 318-329, doi:10.1093/molbev/msw218 (2017).

13 Fadhlaoui-Zid, K. et al. Genome-wide and paternal diversity reveal a recent origin of human populations in North Africa. PLoS One 8, e80293, doi:10.1371/journal.pone.0080293 (2013).

14 Gallego Llorente, M. et al. Ancient Ethiopian genome reveals extensive Eurasian admixture throughout the African continent. Science 350, 820-822, doi:10.1126/science.aad2879 (2015).

15 Schuenemann, V. J. et al. Ancient Egyptian mummy genomes suggest an increase of Sub-Saharan African ancestry in post-Roman periods. Nat Commun 8, 15694, doi: 10.1038/ncomms15694 (2017).

16 Rodriguez-Varela, R. et al. Genomic Analyses of Pre-European Conquest Human Remains from the Canary Islands Reveal Close Affinity to Modern North Africans. Curr Biol 27, 3396-3402 e3395, doi:10.1016/j.cub.2017.09.059 (2017).

17 Garcia-Borja, P., Aura-Tortosa, J. E., Bernabeu-Auban, J. & Jorda-Pardo, J. F. Nuevas perspectivas sobre la neolitizacion en la cueva de Nerja (Malaga-España): La ceramica de la sala del vestibulo. Zephyrus LXVI, 109-132 (2010).

18 Pennarun, E. et al. Divorcing the Late Upper Palaeolithic demographic histories of mtDNA haplogroups M1 and U6 in Africa. BMC Evol Biol 12, 234, doi:10.1186/1471-2148-12-234.; eng; ID: 4230 (2012).

19 Secher, B. et al. The history of the North African mitochondrial DNA haplogroup U6 gene flow into the African, Eurasian and American continents. BMC evolutionary biology 14, 109, doi:1471-2148-14-109 [pii] (2014).

20 Haak, W. et al. Ancient DNA from European early neolithic farmers reveals their near eastern affinities. PLoS Biol 8, e1000536 (2010).

21 Alexander, D. H., Novembre, J. & Lange, K. Fast model-based estimation of ancestry in unrelated individuals. Genome research 19, 1655-1664, doi: 10.1101/gr.094052.109 [doi] (2009).

22 Maca-Meyer, N. et al. Ancient mtDNA analysis and the origin of the Guanches. Eur J Hum Genet 12, 155-162 (2004).

23 Patterson, N. et al. Ancient Admixture in Human History. Genetics 192, 1065-+, doi:10.1534/genetics.112.145037 (2012).

24 Lazaridis, I. et al. Ancient human genomes suggest three ancestral populations for present-day Europeans. Nature 513, 409-413, doi:10.1038/nature13673 (2014).

25 Sanchez-Quinto, F. et al. North African populations carry the signature of admixture with Neandertals. PLoS One 7, e47765, doi:10.1371/journal.pone.0047765 (2012).

26 Prufer, K. et al. The complete genome sequence of a Neanderthal from the Altai Mountains. Nature 505, 43-49, doi:10.1038/nature12886 (2014).

27 Green, R. E. et al. A draft sequence of the Neandertal genome. Science 328, 710-722 (2010).

28 Cortes Sanchez, M. et al. The Mesolithic-Neolithic transition in southern Iberia. Quaternary Research 77, 221-234 (2012).

29 Carpenter, M. L. et al. Pulling out the 1%: Whole-Genome Capture for the Targeted Enrichment of Ancient DNA Sequencing Libraries. American Journal of Human Genetics 93, 852-864, doi:10.1016/j.ajhg.2013.10.002 (2013).

30 Lindgreen, S. AdapterRemoval: easy cleaning of next-generation sequencing reads. BMC research notes 5, 337, doi:10.1186/1756-0500-5-337 (2012).

31 Li, H. & Durbin, R.Fast and accurate short read alignment with Burrows-Wheeler transform. Bioinformatics 25, 1754-1760, doi:10.1093/bioinformatics/btp324 (2009).

32 Li, H. et al. The Sequence Alignment/Map format and SAMtools. Bioinformatics 25, 2078-2079, doi:10.1093/bioinformatics/btp352 (2009).

33 Ginolhac, A., Rasmussen, M., Gilbert, M. T., Willerslev, E. & Orlando, L. mapDamage: testing for damage patterns in ancient DNA sequences. Bioinformatics 27, 2153-2155, doi:10.1093/bioinformatics/btr347 (2011).

34 Skoglund, P., Stora, J., Gotherstrom, A. & Jakobsson, M. Accurate sex identification of ancient human remains using DNA shotgun sequencing. Journal of Archaeological Science 40, 4477-4482 (2013).

35 Kloss-Brandstatter, A. et al. HaploGrep: a fast and reliable algorithm for automatic classification of mitochondrial DNA haplogroups. Hum Mutat 32, 25-32, doi:10.1002/humu.21382 (2011).

36 Schroeder, H. et al. Genome-wide ancestry of 17th-century enslaved Africans from the Caribbean. Proc Natl Acad Sci U S A 112, 3669-3673, doi: 10.1073/pnas.1421784112 (2015).

37 Price, A. L. et al. Principal components analysis corrects for stratification in genome-wide association studies. Nat Genet 38, 904-909, doi:10.1038/ng1847 (2006).

38 Wang, C. et al. Ancestry estimation and control of population stratification for sequence-based association studies. Nat Genet 46, 409-415, doi:10.1038/ng.2924 (2014).

39 Purcell, S. et al. PLINK: A tool set for whole-genome association and population-based linkage analyses. American Journal of Human Genetics 81, 559-575, doi:10.1086/519795 (2007).

40 A language and environment for statistical computing (R Foundation for Statistical Computing. Vienna, Austria. ISBN 3-900051-07-0, URL http://www.R-proiect.org, 2008).

